# Light Amplification By Biofilm And Its Polarization Dependence

**DOI:** 10.1101/139741

**Authors:** Sanhita Ray, Anjan Kr Dasgupta

## Abstract

We report amplified, transmitted light intensity, compared to input, when photosynthetic biofilms were placed in the path of Rayleigh scattered, monochromatic light. Enhancement spectrum shows peak at around 505 nm, which corresponds to the pore wall thickness in biofilm ultra-structure, suggesting role of resonant Mie scattering. Enhancement factors differed when biofilms from different stages of growth were used. Enhancement factors were found to depend on the nature of Rayleigh scattering liquid. Polarizing Rayleigh scattered light by the use of polarizers affected the percentage of enhancement. Amplified output is achievable with constructive interference arising out of coherent forward light scattering, a theoretically predicted outcome of Anderson localization of photons. Possible uses of photosynthetic biofilms in organic material based photonic devices have been discussed.

## 1 Introduction

Biofilms are well-organized aggregates of microbial cells found in diverse habitats. Some of the most functionally important biofilms are photosynthetic since they provide pointers for efficient solar energy harvesting(1). Microstructures of biofilms have been found to be optimized (2, 3) for light capture functions. A mentionable attribute valid for a wide variety of biofilms is the fractal distribution of their component cells (4). Therefore it needs to be noted that biofilms may be looked upon as a 3-dimensionally organized distribution of Mie scatterers (5,6), with scatterer dimension comparable to wavelength of incident beam.

Our attempted investigation of transmittance of light through biofilms revealed some anomalous aspect of light intensity in the forward scattering (FS) direction. Usually a decrease of light intensity (at δ λ = 0) is expected if a media is placed in a light path (unless it is a gain medium as in random lasing). The fact that optical properties of biofilms have special status owing to the presence of dense matrix of exopolymers has been pointed out by earlier workers (7–9). The earliest mention that photosynthetic biofilms can have special optical properties was fron Losee and Wezel (10). However most literature, is regarding the light attenuation through such biofilms. The other intriguing aspect like wavelength (11) and/or polarization dependent amplification of scattered light (12) have got little attention in earlier literature.

The present paper reports for the first time, enhanced light transmission, compared to input, through de-pigmented biofilm of photosynthetic bacteria. The mechanism of such enhanced transmittance, is described in terms of coherent forward scattering, which has been theoretically predicted for strong Anderson localization (13, 14). Rayleigh scattered input has been observed to be an important determinant in such enhancement.

## 2 Results

Biofilms of photosynthetic bacteria *Rhodobacter capsulatus* were grown on both sides of glass cover slip (see Methods) and treated with methanol in order to extract all coloured pigment. The samples were thoroughly dried and placed in the path of Rayleigh scattered light towards photon detector (PD). Elastically scattered irradiation from biofilms was measured, and compared to Rayleigh intensity in absence of biofilm.

In the set-up shown in Figure (1a), initial irridation *(Ir)* from fluorimeter lamp undergoes Rayleigh scattering by milliQ water in the cuvette. Input (*I*) to biofilm was taken as the Rayleigh intensity measured in the direction of photodetector (PD, placed at 90 deg w.r.t. *Ir*) with empty slit (S, figure 1(b)) being placed in the light path. The fraction of the Rayleigh scattered (*R*) light that passes through the empty slit forms the incident irradiation on the biofilm. Output (*O*) was taken as the total light that passed through the slit and was detected by the PD, when biofilm was mounted behind S i.e. placed in the light path of *I*. Figure (1(c)) shows that the total transmitted output (*O*) was greater than incident intensity (*I*, given as “blank” in all figures) i.e. *O > I*, and the change was much greater than PD noise. The times enhancement (XI) i.e. O:I ratio calculated from this data is shown in figure 1(d). Placing empty glass cover-slip behind the slit resulted in light absorbance upto 390 nm and intensities remained same as blank at higher wavelengths. Enhancement increased with increasing wavelength, but showed peaks at 505 nm and a less distinct peak at around 630 nm.

**Figure 1:**
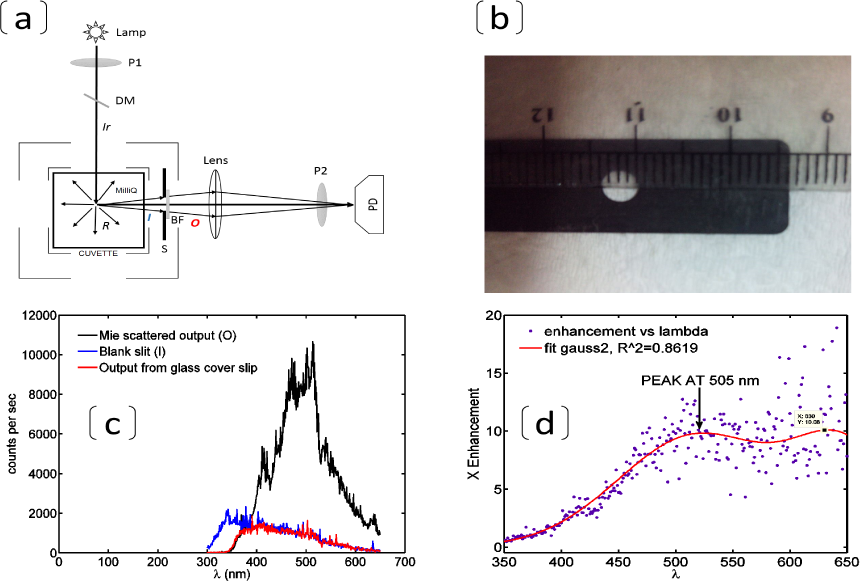
(a) Shows the set-up used for observing transmitted light through biofilm. DM= dichroic mirror, R=Rayleigh scattered light, I=input irradiation to biofilm, O=output light from biofilm, Ir=excitation beam from fluorimeter lamp, P1=excitation polarizer, S=slit, BF=biofilm, P2=emission polarizer, PD=photon detector. (b) Slit (diameter=4 mm) placed in light-path between cuvette and PD, either empty or with biofilm. (c)Shows input inout intensity when slit was blank, output obtained from alcohol treated biofilm placed on slit and intensity obtained when empty glass cover-slip was placed on slit. Shows times enhancement i.e. output:input ratio as a function of wavelength and fitted to a double-gaussian curve. Two peaks were obtained, at 505 nm and at 630 nm.

Light source was a non-coherent, unpolarized beam from fluorimeter lamp. *Ir* was made monochromatic by the use of excitation monochromator. This was used in conjunction with an emission monochromator, such that, at each wavelength during scanning, emission was detected at the same wavelength as excitation (15) (see Methods). This ensures that only elastically scattered photons are detected.

The experimental set-up directly measures flux through a 4 mm diameter circular slit (see figure 1(b)) in the forward direction, as collected by the lens placed 3.5 cm in front of the slit. Forward direction is with respect to direction of propagation of Rayleigh scattered light through the slit, towards detector. It should be noted that two seperate elastic scattering events are contributing to the results, at two seperate sites:

- Rayleigh scattering inside cuvette (since water molecules are much smaller compared to the wavelength of incident light): measured with blank slit.
- Mie scattering within porous biofilm (since the dimension of pore walls are comparable to wavelength of visible light, see additional data, Figure S1: output of such scattering was measured when biofilm was placed in the path of Rayleigh input.

Placement of a translucent barrier in the path of such Rayleigh scattered light is expected to decrease the detected intensity. Obtained enhancement may be minimally explained on the basis of constructive interference (CI) of photons. The porous ultrastructure of RCSB biofilm (as revealed by SEM imaging, see additional data, Figure S1) may be considered as a 3-dimensional fractal network (4) of Young slits (16). Incident light may undergo multiple scattering and photons through optical slits interfere constructively, provided they are phase matching. Multiple scattering in disordered media (17) has been theoretically and experimentally studied for photonic localization which gives rise to peak in backscattered intensity (11, 14, 18–25), which is well known and experimentally studied. More recently, coherent forward scattering (CFS) (13, 15, 26–29) has been predicted for cases of strong photonic localization. Obtained enhancement in forward direction may be explained as occuring due to CFS.

Observed wavelength dependence is consistent with photonic localization since the extent of localization is dependent on wavelength. Wavelength dependence of times enhancement as obtained in our experiments may be explained on the basis of kl* dependence of localization, where *k* = 2 π/λ and l*=mean free path. For strong localization the value of kl* should be as near to 1 as possible. Hence, extent of localization should increase as λ increases, thereby bringing the value of kl* near unity. For our case, considering average pore diameter to be the mean free path, l*=200 nm and therefore kl*=2.5 at 500 nm. At 600 nm, the value of kl*=2.0944. Enhancement spectra showed peak at 505 nm. This corresponds to average lateral dimension of the connected cellular structure ( see additional data, Figure S1) and hence points towards resonant Mie side scattering.

### 2.1. Dependence on biofilm structure

Times enhancement with varying samples of biofilm was investigated by using biofilms collected on 4th, 5th, 6th and 7th days of growth and treated with 100% alcohol (see figure 2(a)). From 4th day onwards times enhancement decreases with days of growth. To account for structural disordering, 7th day film samples were seperately treated with two concentrations of alkali solutions (0.085 M and 0.17 M NaOH) and methanol (see figure 2(b)). It was previously observed that strong alkali solutions disrupt biofilms significantly whereas nearly all other reagents were seen to leave the architecture intact. Treatment with the stronger alkali solution resulted in increased enhancement compared to the weaker one which was approximately the same as with methanol treatment. The results indicate that lesser thickness probably contributes to the emergence of enhanced output. Alternatively, greater thickness leads to absorption or complete inhibition of transport by Anderson localization.

**Figure 2:**
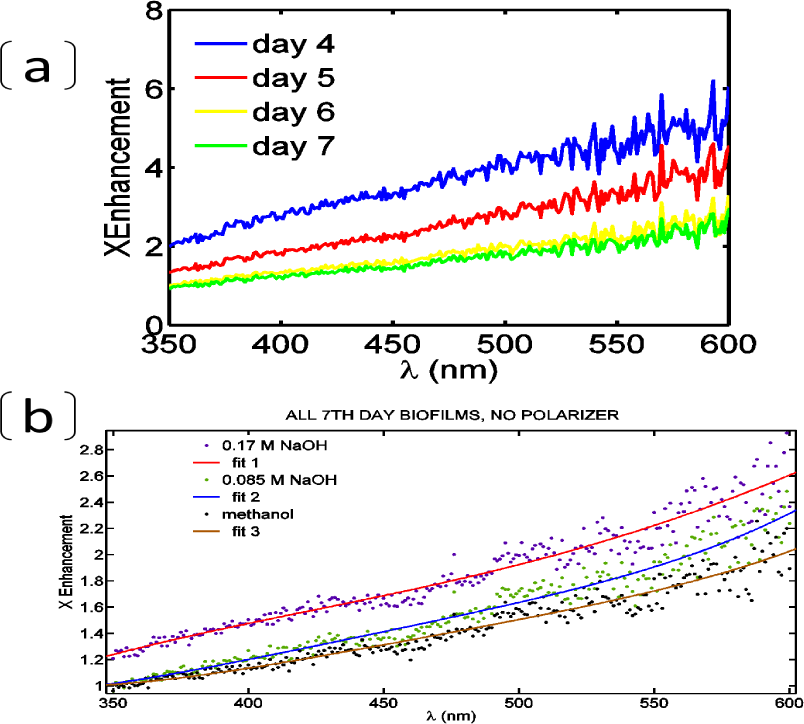
(a) Times enhancement as a function of wavelength (*XI*(λ)) are compared for four biofilm samples collected on 4th, 5th, 6th and 7th days (COLOR online) of biofilm growth and treated with methanol. All showed increase of XI with λ but at any particular value of λ XI decreased with more number of days of growth. (b) *XI*(λ) for replicate 7th day biofilm samples, treated with either 0.17 M or 0.085 M NaOH solutions (known as disruptive agents for biofilms), are compared with that methanol treated 7th day biofilm. Note that biofilm residues were still present after alkali treatment though pigmentation was lost. Higher concentration of disruptive agent resulted in increased enhancement, whereas lower concentration was almost same as methanol treated sample.

Decrease in mean free path has been reported to decrease the extent of localization as shown by Etemad et al ((30)) The decrease in enhancement values with more days of growth (figure 2(b)) probably reflects this in terms of more packed (hence lesser mean free path) cellular distribution. This result is further supported by the fact that treatment with higher concentration of alkali results in increased enhancement (figure 2b) compared to lesser concentration. These results effectively demonstrate that the enhancement phenomenon is dependent on biofilm.

### 2.2 Liquid medium dependence

Enhancement was investigated by putting various liquids in the cuvette (see figure 3 (e)), instead of water, like acetone and glycerol. Water showed the highest times enhancement followed by glycerol, with acetone showing lowest enhancement. The order of increase in enhancement factor may be seen to follow the order of increasing dielectric constant, as well as relative polarity. This suggests a role of solvent in tuning spatial or temporal coherence of Rayleigh scattered light that is incident on the biofilm.

**Figure 3:**
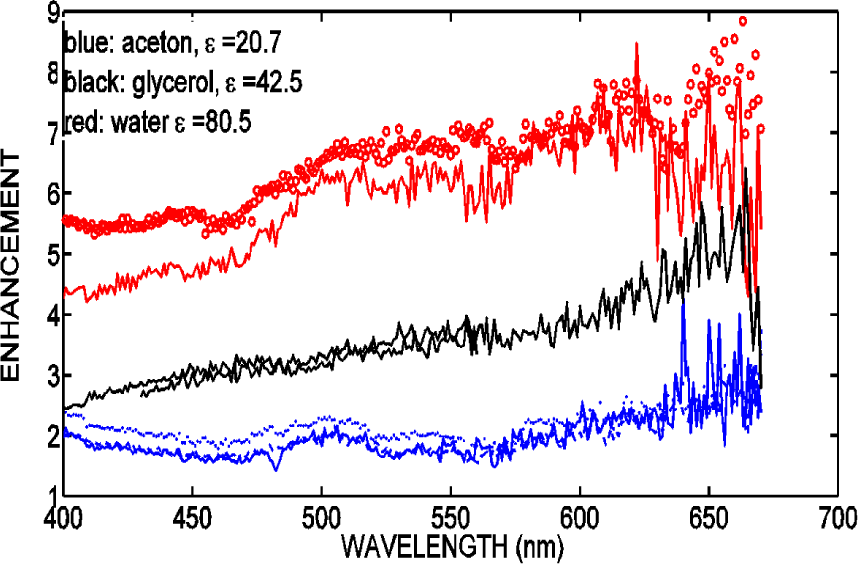
Effect of varying the liquid medium for Rayleigh scattering. Acetone and glycerol were taken in cuvette as Rayleigh scattering medium and this was observed to affect the enhancement factor.

### 2.3 Polarization Effects

In order to investigate role of spatial coherence, input light to biofilm was polarized by the use of excitation polarizer (P1, that polarizes *Ir),* see figure 4(A and B) as well as a sheet polarizer (SH, that polarizes *I*), see figure 4 (E). Polarization analysis was done by placing an analyzer polarizer immediately before photodetector. All polarizers can be rotated to obtain rotation of their plane of polarization.

**Figure 4:**
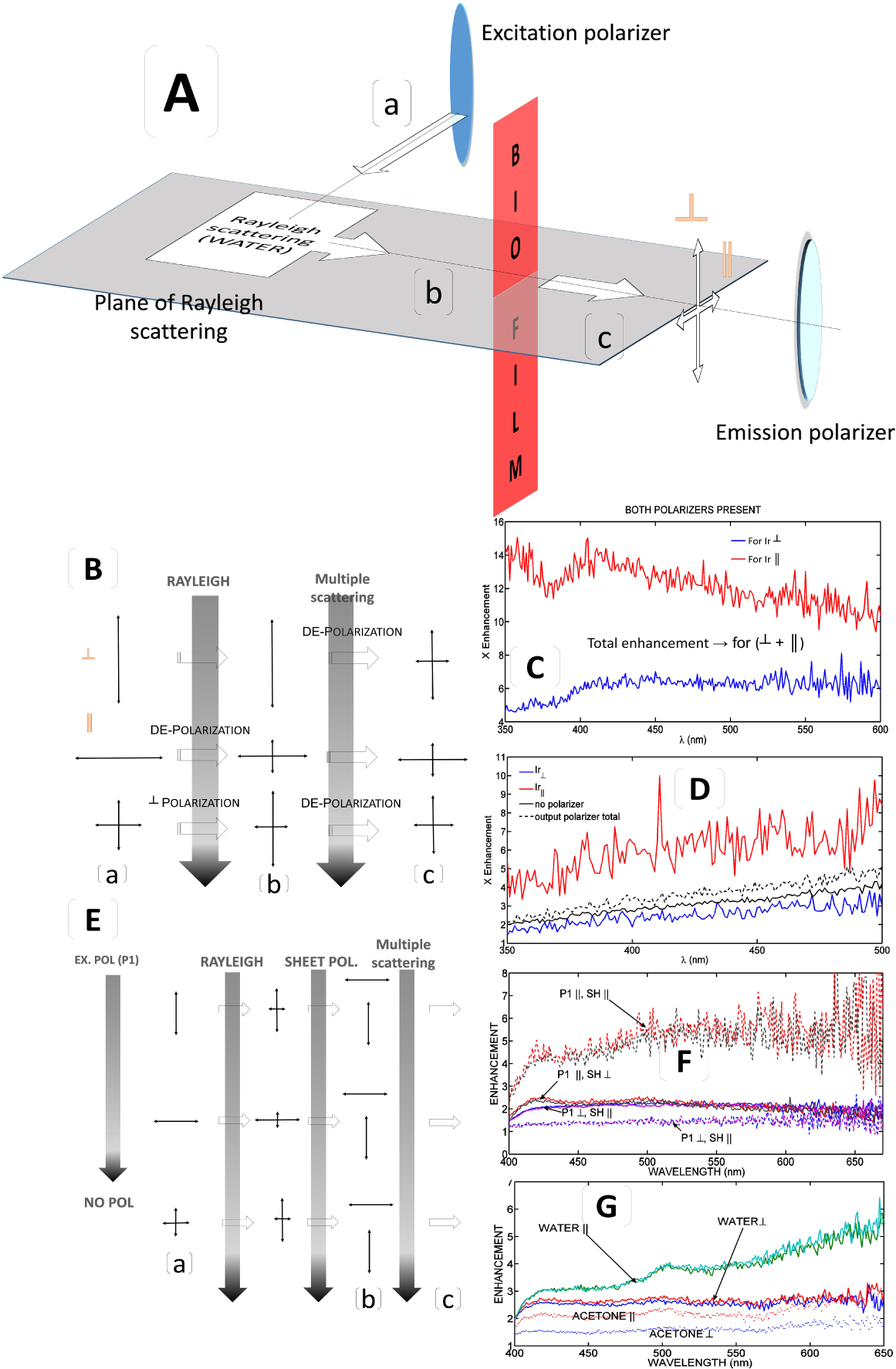
Effect of polarizing input light on enhancement factor (A) Plane of Rayleigh scattering is shown and perpendicular (⟘) and parallel (||) alignments defined with respect to this plane. (B) Polarization states (a) before Rayleigh (b) after Rayleigh and (c) after multiple scattering illustrated using arrow diagrams. (C) Effect of *Ir* polarization (by use of P1, excitation polarizer) on enhancement, calculated by summing both parallel and perpendicular contribution, as detected by the use of P2 (emission polarizer). (D) Times enhancement, using only excitation polarizer, are shown when its plane of polarization is parallel and perpendicular to the plane of Rayleigh scattering, and compared to when there was no excitation polarizer and when there was only emission polarizer. Other than the last case, no emission polarizer was present. (E) Polarization states (a) before Rayleigh and (b) after passing through sheet polarizer illustrated using arrow diagrams. (F) Effect of using combination of excitation polarizer and sheet polarizer, to completely polarize incident radiation on biofilm. Note that sheet polarizer is placed before the biofilm, on the side of the cuvette. (G) Effect of using only sheet polarizer, in different alignments,on enhancement. Enhancement factors are also for similar experiments, when acetone is the scattering liquid.

Excitation polarizer P1 was placed in the path of irradiation from lamp (*Ir*), keeping plane of polarization either vertical or horizontal. Similarly, analyzer polarizer P2 was placed immediately before the photon detector. Linear polarization of detected radiation (either Rayleigh through blank slit or scattered output from biofilm on slit) was determined by keeping its plane of polarization along vertical or horizontal. The vertical direction is designated at all places as perpendicular, ⟘, i.e. perpendicular with respect to plane of Rayleigh scattering in experimental set-up. The horizontal direction is designated at all places as parallel, ||, i.e. parallel with respect to plane of Rayleigh scattering.

Consequently *Ir_⟘_* and *Ir*_||_ denote irradiation from fluorimeter lamp with plane of polarization along vertical and horizontal directions respectively. *I_⟘_* and *Ir*_||_ denote detected intensities from blank slit keeping plane of P2 along vertical and horizontal respectively. *O_⟘_* and *O*_||_ denote detected intensities from biofilm-mounted slit keeping plane of P2 along vertical and horizontal respectively. *XI ⟘* denotes times enhancement obtained with *Ir_⟘_*. *XI*_||_ denotes times enhancement obtained with Ir_||_.

Enhancements were calculated seperately for when *Ir_⟘_* and *Ir*_||_ were used. In each case, enhancement was calculated taking into account *total O,* i.e. summation of parallel and perpendicular components of Mie scattering from biofilms (see Methods, available online), and its ratio to *total I* i.e. summation of parallel and perpendicular components of Rayleigh scattering. Please note these calculations are for cases where both polarizers were used simultaneously.

Higher percentage enhancement was obtained (see figure 4 (C)) with parallel polarization of incident irradiation (*Ir*_||_) from lamp, compared to perpendicularly polarized irradiation *(Ir_⟘_*). We may correlate the percent enhancement to polarization of Rayleigh scattered radiation (see additional data, Figure S3) in each case. Thus, vertically polarized Rayleigh scattered input is shown as giving rise to lower enhancement, compared to partially horizontally polarized Rayleigh input (for polarization values along the entire spectrum, see additional data, Figure S3).

Difference of enhancement with *Ir_⟘_* and *Ir*_||_ prompted the authors to investigate the effect using only excitation polarizer P1-either with plane of polarization in the vertical direction (i.e. perpendicular to the plane of Rayleigh scattering) or in the horizontal direction (i.e. parallel to the plane of Rayleigh scattering). Note that no emission polarizer P2 was used for this measurement. Times enhancement was calculated as *XI_⟘_ = O|Ir_⟘_|I|Ir_⟘_* and *XI*_||_ = *O\Ir*_||_*)/(I_⟘_*) for *Ir*_||_ and *Ir_⟘_* respectively. Enhancement (see figure 4(D)) with vertically polarized incident radiation *(Ir_⟘_)* i.e. was lesser than that with horizontally polarized incident radiation (*Ir*_||_) i.e. *O_⟘_: I*_||_ *> O*_||_: *I*_||_. Both are shown compared to when no polarizer was used, which was slightly greater than *Ir_⟘_.*

Role of Rayleigh input on enhancement was investigated by placing a sheet polarizer immediately before the biofilm, and rotating its plane of polarization (figure 4(A, E-G)). This results in polarization based filtering of Rayleigh scattered light and chosen polarization (for polarization values along the entire spectrum, see additional data, Figure S5) was exclusively incident on the biofilm. It was found that even in this configuration, horizontal polarization produced greater enhancement compared to vertical polarization (figure 4(G)). It should be mentioned here that biofilms are quasi-2-dimensional thick films and in the measurement set-up the plane of biofilms is oriented at right-angles to the direction of propagation of light from cuvette to detector. Hence there is no chance of horizontal direction being preferred to vertical direction, in the current set-up, see figure 4(A).

The only specified plane in the experimental set-up is the plane of Rayleigh scattering. Hence all polarization effects should trace back to Rayleigh scattering and how it affects the quality of input to biofilm. Excitation polarizer was used in conjunction with sheet polarizer, see figure 4 (E). Parallel versus perpendicular alignment of excitation polarizer did not affect enhancement significantly when sheet polarizer was vertically oriented. However, when sheet polarizer had its plane of polarization parallel to plane of Rayleigh scattering the percentage enhancement was greater, see figure 4 (F).

It becomes clear from the results that enhancement is polarization encoded and hence controlled by Rayleigh scattering. For application purposes, we may tune enhancement signal by modulating the polarization of the input. Alternatively, for thick film characterization, polarization indices may become a distinguishing parameter. Additionally, the (Mie) output polarization was found to depend on input conditions (see additional data, Figures S4 and S5) and may be utilized for such multi-parametric determination.

## Implications

### 3.1 *Non-coherence* vs *coherence induction*

All previous experiments on localization have used phase coherent light source. In our case, Rayleigh scattered light was used. Rayleigh scattering affects both the phase and polarization of incident radiation. Additionally the light source used was fluorimeter lamp with no spatial or temporal coherence. These factors may be responsible for CFS effects being observable in macroscopic time. Effect of changing the liquid medium and input polarization supports a strong role of Rayleigh scattering in bringing about the light amplification. It is to be noted that the amplification factor increases in an opposite manner to Rayleigh intensity over the wavelength spectrum studied.

### 3.2 Ecological importance

It should be pointed out that all previous localization experiments used coherent laser source whereas ours was a non-coherent source. A detectable enhancement, with non-coherent source, points to biofilm architecture as a coherence inducing media. Thus we may consider interior photosynthetic layers as well as more bottom dwelling species as receiving phase coherent light. This points to biofilms having evolved a micro-structure that achieves coherence and other optical tuning. This is of great importance considering the previous literature on entanglement phenomenon (31) in photosynthesis.

Most of the measurements of photo-synthetically important microbial mats have been measured using scalar irradiance and spectral attenuation (32). The point that has been often overlooked in such measurements is the Mie-resonant nature of the scattering that provides optimal peak of such irradiance at resonant wavelength. The obtained peaks were correlated with absorption maxima of pigments. We circumvent the absorption problem by treating photosynthetic biofilms with methanol and extracting pigments. Our results clearly show that in the absence of pigments, enhancement peaks are shown, resonant peaks being a typical signature of Mie scattering. The way in which such biofilm will scatter (33) the incident radiation to its local microbial community is evidently of ecological significance. Literature demonstrates the role of symbiotic hosts of photosynthetic biofilms.

Any light enhancing property observed for structured ensemble of photosynthetic organisms points towards ecological importance of such properties in the niche it occupies (see additional data, Figure S6). Purple non-sulphur bacteria are benthic organisms living underneath a layer of plankton and aerobic benthos. As such sunlight reaching deep benthic layers is Rayleigh scattered output from the upper layers, having an approximate resemblance to our experimental set-up. RCSB forms thick mats and hence the enhanced output is probably available to interior cells. Rayleigh scattering decreases with increase in wavelength and enhanced output at higher wavelengths probably helps to inverse this deficit. One importance of the wavelength dependence may be to cut out on high energy irradiation in the UV/blue range.

Enhanced scattering (compared to incidence) has been previously reported for multiple scattering by spicules within tunic layers of ascidians (9, 32–34) that host *Prochloron* (cyanobacteria) as obligate symbionts. However the authors have not shed any light on this enhancement phenomenon. Their data shows λ dependence similar to our case. Incidentally, this enhanced light is available for photosynthesis by the obligate symbionts. There remains the possibility of some of the troughs corresponding to Mie peaks as opposed to absorbance. They have conducted their studies using live, pigment-containing biofilms which would have masked any enhancement taking place within cyanobacterial biofilms.

### 3.3 Possible application areas

Random lasers (21, 35) consist of gain media that function on the basis of constructive interference that results from multiple scattering. Optimizing random gain media is a challenge. Structures like biofilms may be utilized for constructing soft material based random lasers by using a trapped film of Rayleigh scattering liquid. Biofilm residues can be utilized for lesser energy consuming, smart lighting solutions, as coating layer for LEDS. Light capture efficiency is a primary target for improving solar cells-de-pigmented biofilm materials may be used as coating to improve energy yield, as in case of of living photosynthetic bacteria. Anderson localization has been utilized for photonic signal transmission through optical fibres (21, 36). The polarization encoding (see supplementry figures S3–S5) that is demonstrated in our amplification studies, suggests biofilms may be used for signal transmission. It is possible to grow biofilms in or on capillary tubes or fibers. Finally, biofilms are a major bio-medical and industrial concern and this technique may be used as a characterization tool for biofilms (5, 37) as well as other kinds of thick films.

## 4 Conclusion

In this work, intensity of forward scattered light from biofilms has been shown to be greater than incident light intensity, at the same wavelength. Enhancement spectrum showed possible resonance peaks. The phenomenon has been explained on the basis of constructive interference as a result of coherent forward scattering. The extent of intensity amplification has been shown to be sensitive to variations in the biofilm. Our findings present photosynthetic biofilms as a coherent barrier to incoming, randomly scattered, light. The extent of amplification shows clear polarization encoding. In conclusion, the authors would like to note that this is very much a work in progress towards determining how Rayleigh scattering contributes towards enhanced output, as well as the role of polarization in controlling the extent of enhancement.

## 5 Supplementary

Polarization of *I* (from Rayleigh scattering) and *O* (from biofilm) were determined (Figure 7 (a)) when *Ir _⟘_* was used and compared to polarization values when *Ir_||_* was used. Note that polarization,

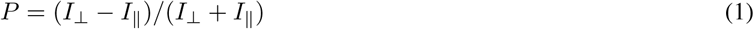

Rayleigh scattered light (*I*) was vertically polarized (P > 0) when *Ir _⟘_* was used and percentage polarization was high (|P|_⟘_ 1). Polarization was partially reversed (P < 0) when *Ir_||_* was used but |*P*|_||_ was lower compared to previous case. Rayleigh scattered light is known to exhibit polarization along perpendicular to plane of scattering and this polarization is highest for 90 deg with respect to incident illumination. This expains the polarization reversal. Mie scattered output (*O*) from biofilms was found to be significantly depolarized (see histogram, 4 (a)), compared to Rayleigh scattered input, when *Ir _⟘_* was used. Change in polarization was almost negligible when *Ir_||_* was used.

Figure 8 shows polarization of *I* compared to *O*, obtained by using only emission polarizer (P2). The Mie scattered output was depolarized compared to input Rayleigh scattered light. This result was in keeping with that of Figure 7. The comparison of enhancement obtained over all wavelengths (data over the entire spectra represented as a boxplot) with polarization (median value from the spectra investigated) of respective input shows that enhancement was greater for depolarized output.

## 6 Materials and Methods

### 6.1 Biofilm preparation

*Rhodobacter capsulatus* SB1003 was cultured in RCV medium, in presence of light and micro-aerobic conditions, keeping pieces of glass cover-slips submerged in the culture medium. Biofilms were usually grown for either 12 days (Figure (1), (2)) or 4 days (Figure (3)). After growth, samples were collected and treated with methanol at 50 deg for 2 hours. The cultured biofilms were red upon harvesting. Samples turned white after methanol treatment. Samples were dried at 50 deg after pigment extraction by methanol treatment.

**Figure S1.**
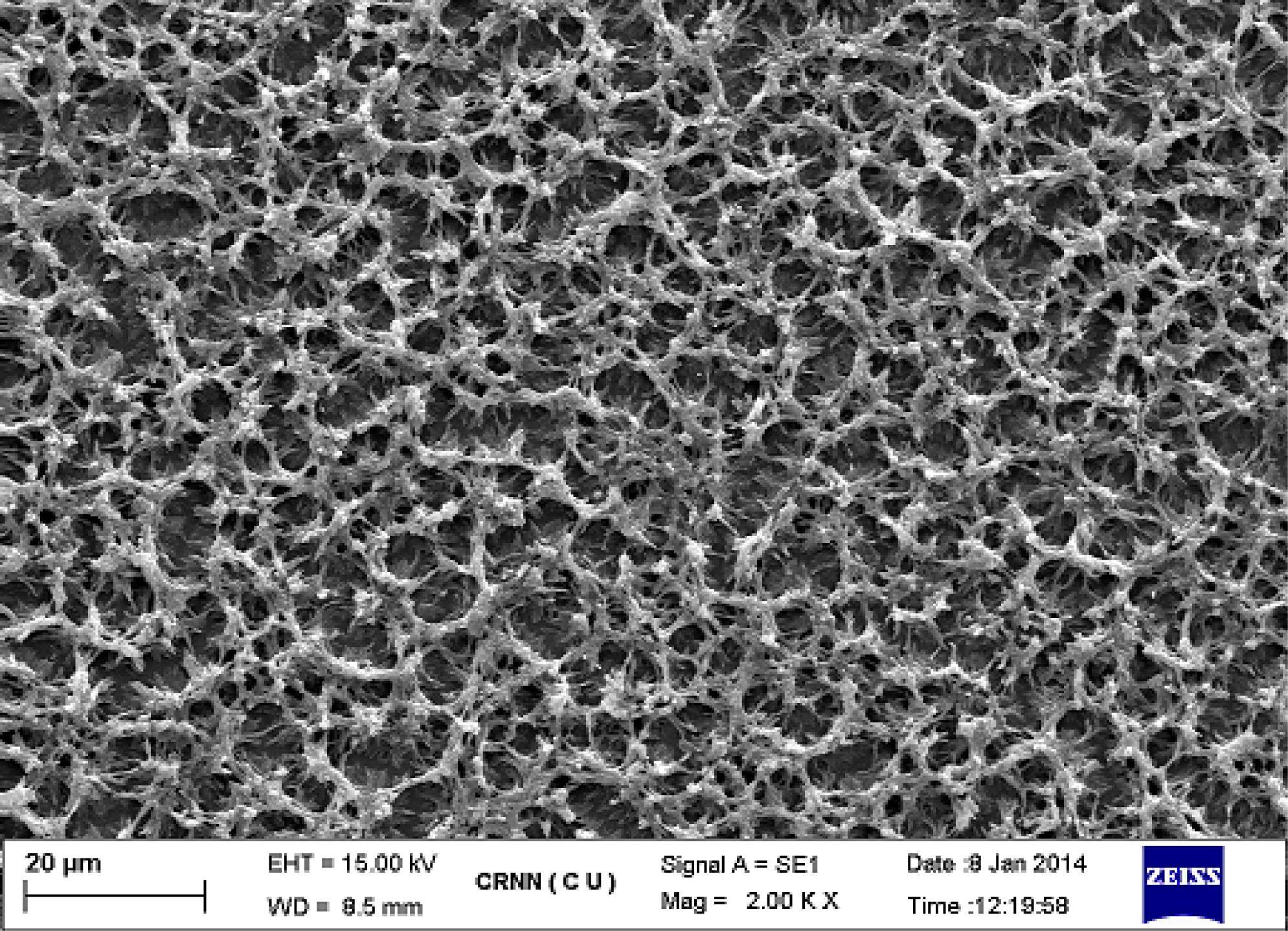
Scanning electron micrograph of 11th day biofilm.

**Figure S2.**
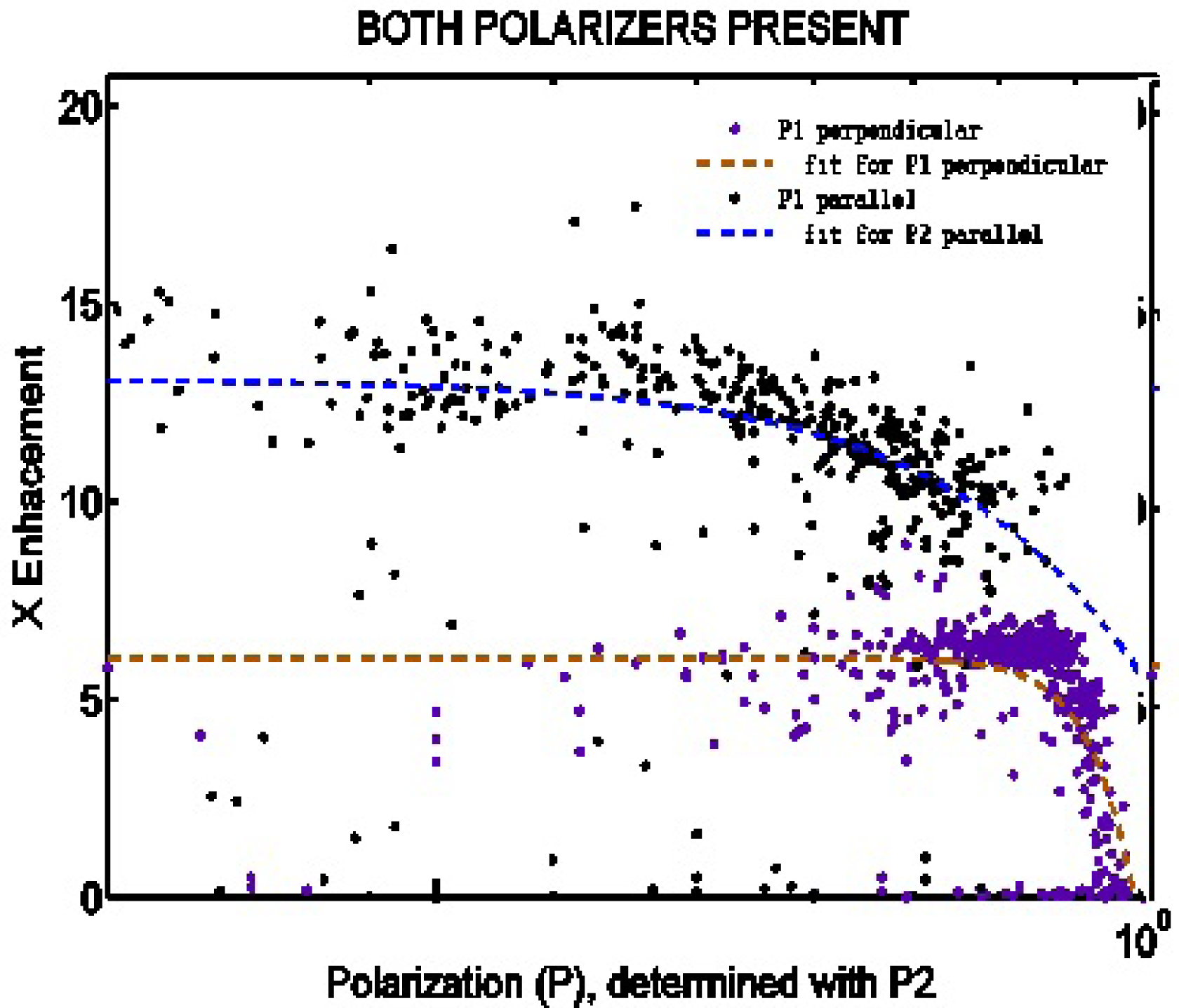
Dependence of times enhancement on absolute values of input polarization (brought about by placing P1 in the path of excitation beam *(Ir)* and due to Rayleigh scattering, detected by placing P2 before photo-detector) was plotted for data collected at various wavelengths, shown in figure (2) (a) and (b), giving polarization and enhancement values respectively. Note that polarization values for horizontal alignment of P1 are negative, here only their absolute values are plotted. For both horizontal and vertical alignments of plane of polarization of P1, times enhancement decreased with absolute values of polarization according to a power function. However for *P*1_||_ values of enhancement were higher than that for *P1*_⟘_ even when absolute values of input polarization were comparable. Note that in the figure x-axis is in the log scale.

**Figure S3:**
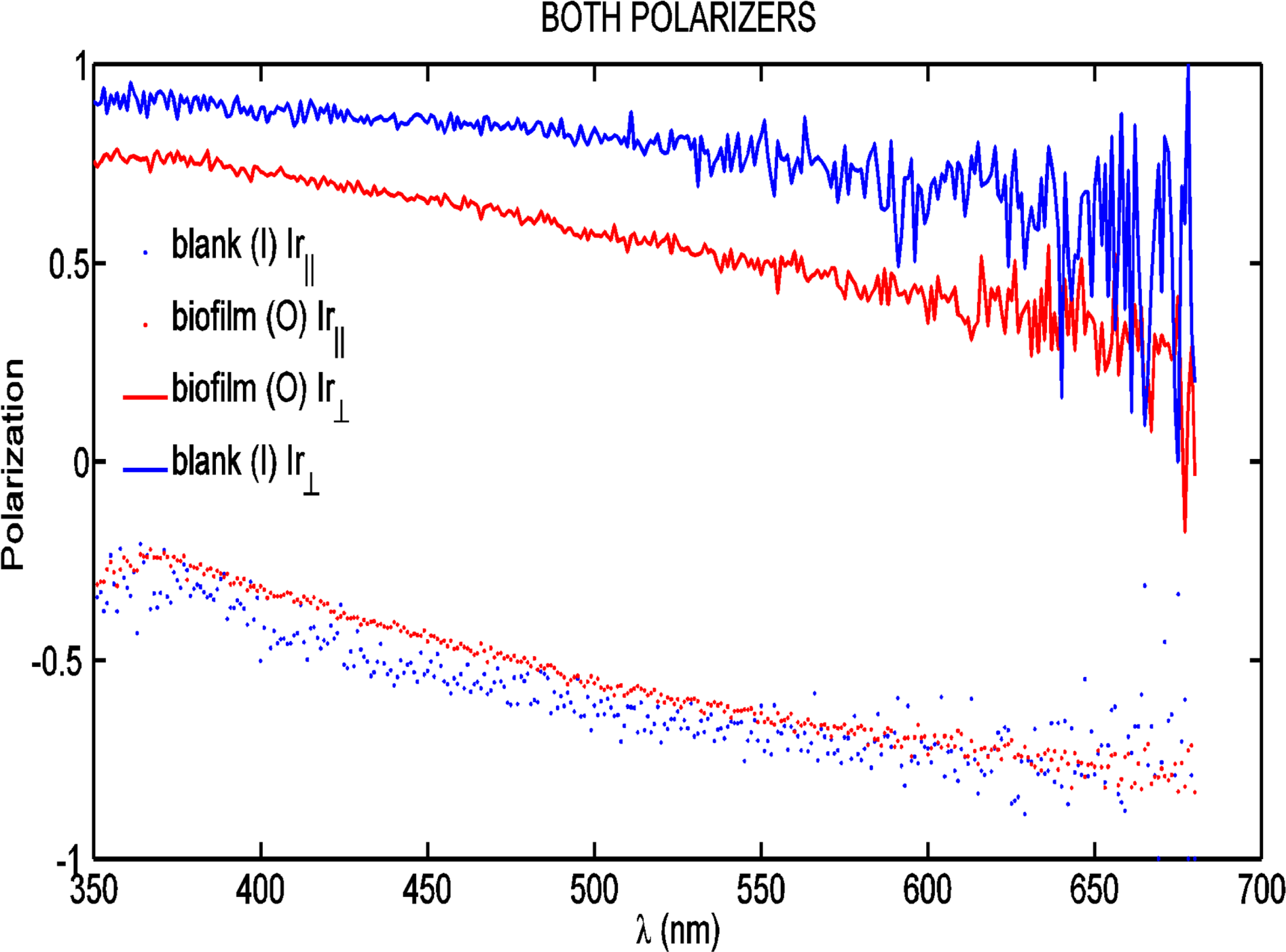
Polarization of input (*I*), denoted as blank, for plane of polarization of P1 along parallel and perpendicular (see text), compared to output (O), denoted as biofilm, for same alignments of P1.

Cover-slips were introduced into 4 seperate tubes containing inoculated media. Cover-slips were collected from each tube on 4th, 5th, 6th and 7th days of growth respectively, treated with methanol and dried. Two more samples of 7th day biofilms were obtained and treated with 0.085 M and 0.17 M NaOH solution at 50 deg, for 2 hours, followed by drying. Treatment with strong alkaline solutions had been previously observed to disintegrate *Rhodobacter capsulatus* biofilms, hence treated samples were expected to have looser cellular packing.

### 6.2 Optical measurements

Light output from biofilms were measured using synchronous fluorescence detection mode in a PTI Quantamaster fluorimeter. Excitation and emission monochromators were set at the same wavelength (*δλ* = 0) for each datapoint i.e. emission was detected at the same wavelength as excitation for each wavelength, thus giving elastically scattered intensities only. A cuvette containing MilliQ water was placed in the standard fluorimetric geometry i.e. with photon detection (PD) at right angles w.r.t. incident beam (Ir) from lamp. An empty slit (S, diameter=4 mm) was placed in the path of Rayleigh scattered light (R) between cuvette and PD. Input (I, counts/s) was measured as intensity that passed through the empty slit. To measure output from biofilm, in the FS direction, biofilm grown on pieces of cover-slip were mounted on the slit with the help of adhesive tape- PD counts in this configuration were taken to be the output. Note that biofilms are anisotropic i.e. arrangement of cells in the x-y plane is different from that in the z-direction. In this case light path from cuvette to PD was along z-axis of the biofilm.

For experiments with polarizers, polarizers were either placed in the path of Ir (P1) or immediately before PD (P2), or both. Note that R and hence I is already vertically polarized i.e. at right angles w.r.t. plane of Rayleigh scattering.

### 6.3 Scanning Electron Microscopy

Biofilm samples were fixed with 2.5% glutaraldehyde, dehydrated in graded alcohol, stained with 0.1% osmium tetroxide and scanning electron microscopy (ZEISS EVO-MA 10) was performed after gold sputtering on the sample.

### 6.4 Calculating enhancement with polarizers P1 and P2

Total enhancement when both polarizers (P1 and P2) were used (see figure 4(A)) was calculated as

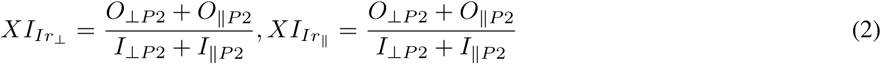

**Figure S4:**
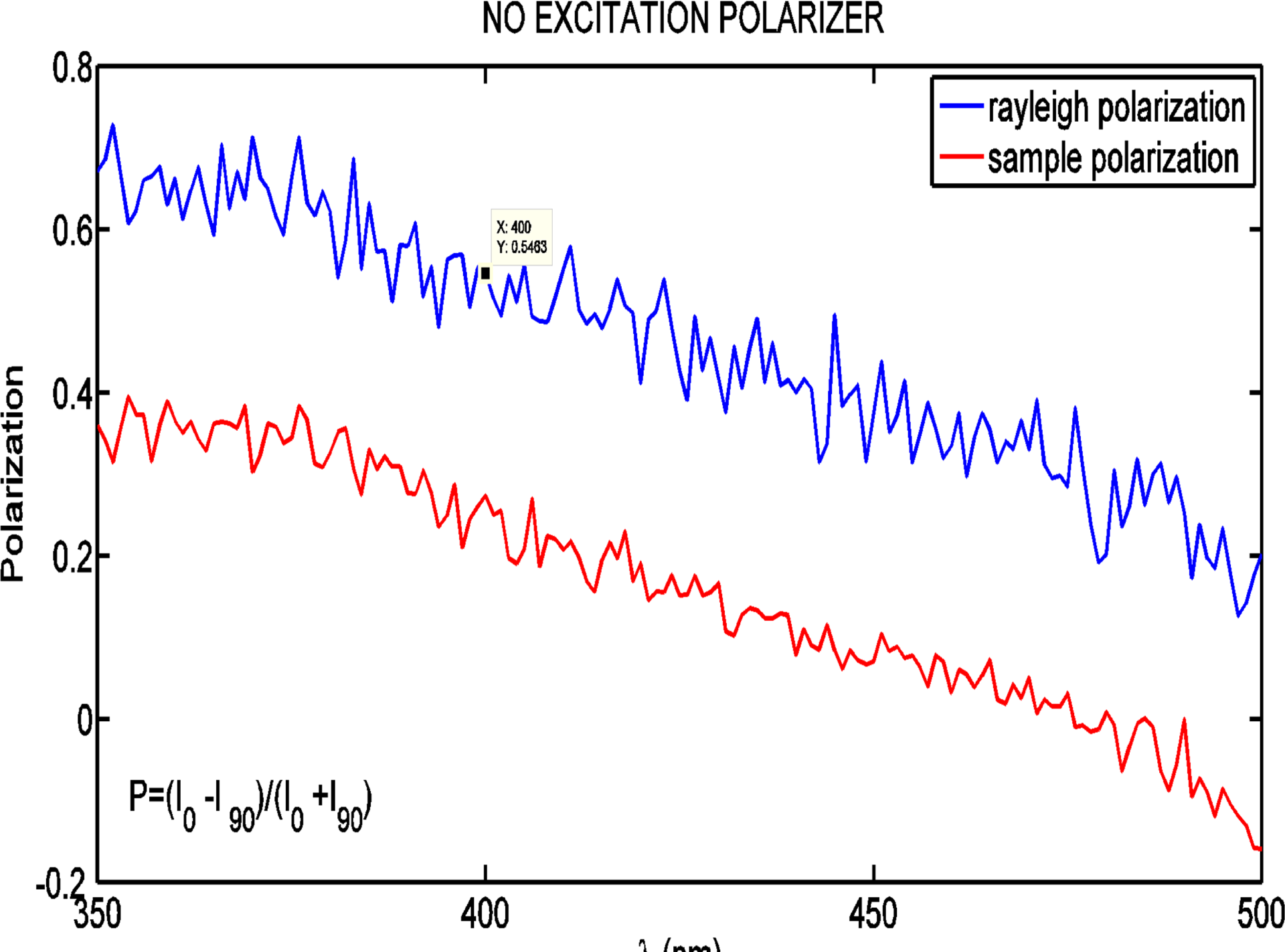
Polarization of input Rayleigh scattered irradiation is shown compared to polarization of output from biofilm. The output is depolarized w.r.t. input. For this experiment, only emission polarizer P2 was used.

**Figure S5:**
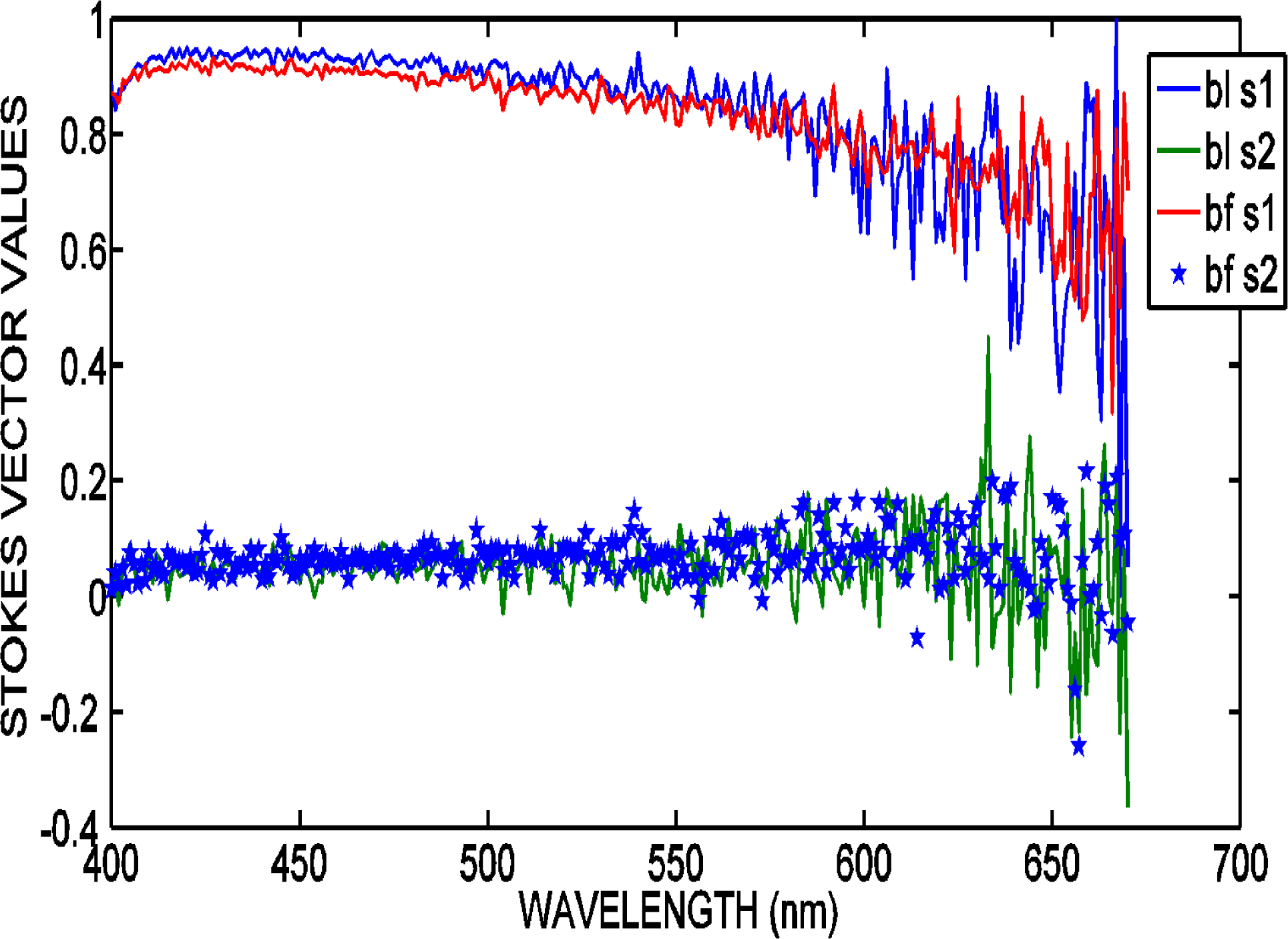
Polarization of Rayleigh scattered irradiation, after passing through sheet polarizer, compared with polarization values for Mie scattering through biofilm, in the same set-up. Sheet polarizer was kept with its plane of polarization perpendicular to the plane of Rayleigh scattering. Unlike in previous cases, no depolarization was observed.

**Figure S6:**
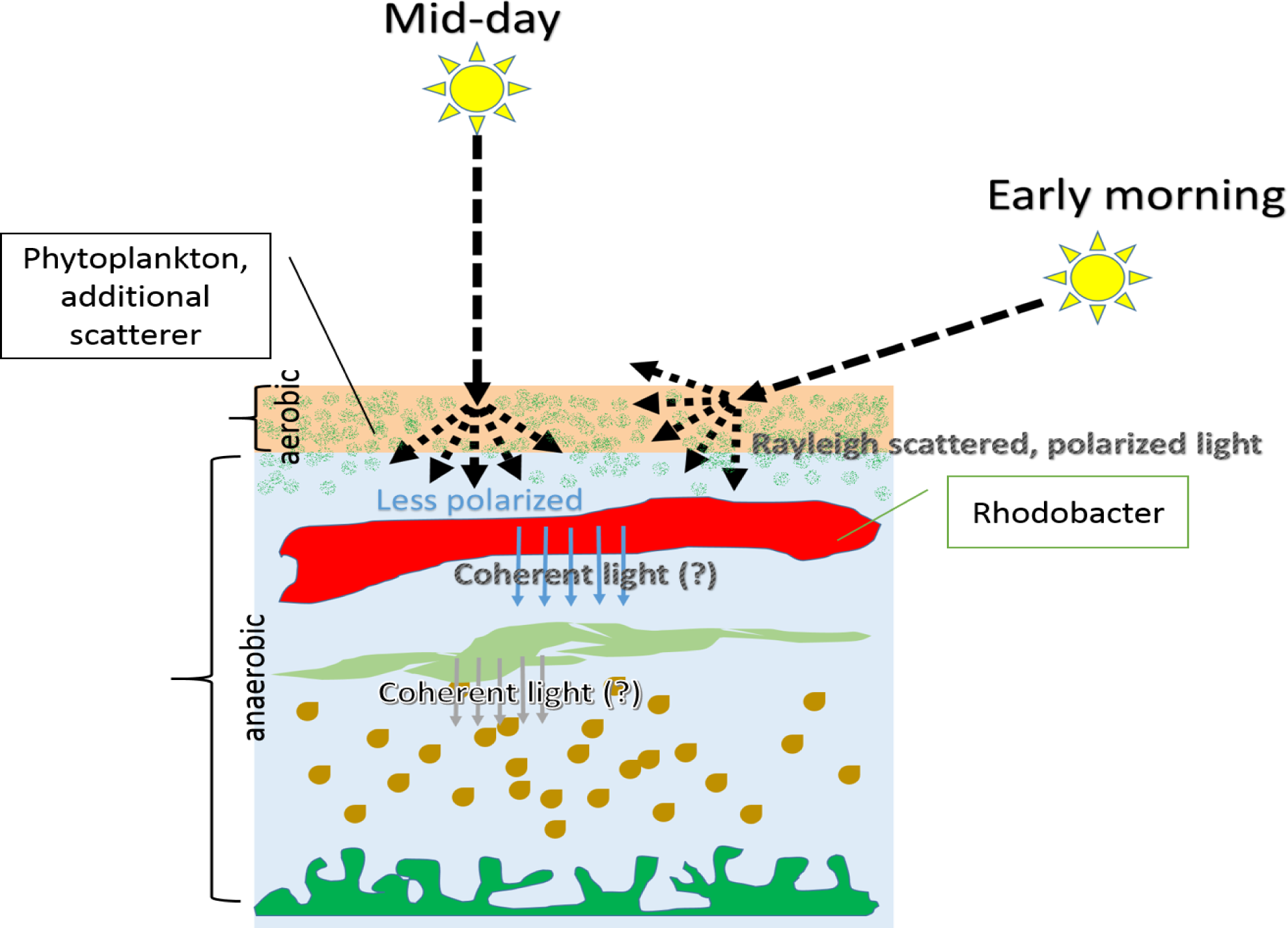
Possible ecological significance of light enhancement and coherence induction on marine niche.

## References

1. Carmeli, I., M. Cohen, O. Heifler, Y. Lilach, Z. Zalevsky, V. Mujica, and S. Richter, 2015. Spatial modulation of light transmission through a single microcavity by coupling of photosynthetic complex excitations to surface plasmons. Nature communications 6.

2. Brodersen, K. E., M. Lichtenberg, P. J. Ralph, M. Kuhl, and D. Wangpraseurt, 2014. Radiative energy budget reveals high photosynthetic efficiency in symbiont-bearing corals. Journal of The Royal Society Interface 11.

3. Domozych, D. S., and C. R. Domozych, 2008. Desmids and biofilms of freshwater wetlands: development and microarchitecture. Microbial Ecology 55:81–93.

4. Hermanowicz, S., U. Schindler, and P. Wilderer, 1995. Fractal structure of biofilms: new tools for investigation of morphology. Water Science and Technology 32:99–105.

5. Beyenal, H., Z. Lewandowski, C. Yakymyshyn, B. Lemley, and J. Wehri, 2000. Fiber-optic microsensors to measure backscattered light intensity in biofilms. Applied optics 39:3408–3412.

6. Cheong, F. C., S. Duarte, S.-H. Lee, and D. G. Grier, 2009. Holographic microrheology of polysaccharides from Streptococcus mutans biofilms. Rheologica Acta 48:109–115.

7. Wilson, B. C., and S. L. Jacques, 1990. Optical reflectance and transmittance of tissues: principles and applications. Quantum Electronics, IEEE Journal of 26:2186–2199.

8. Vogelman, T. C., J. N. Nishio, and W. K. Smith, 1996. Leaves and light capture: light propagation and gradients of carbon fixation within leaves. Trends in Plant Science 1:65–70.

9. Kuhl, M., C. Lassen, and B. Jorgensen, 1994. Light penetration and light-intensity in sandy marine-sediments measured with irradiance and scalar irradiance fiberoptic microprobes Rid A-1977-2009. Marine Ecology-Progress Series.

10. Losee, R. F., and R. G. Wetzel, 1983. Selective light attenuation by the periphyton complex. In Periphyton of Freshwater Ecosystems, Springer, 89–96.

11. Asatryan, A. A., S. A. Gredeskul, L. C. Botten, M. A. Byrne, V. D. Freilikher, I. V. Shadrivov, R. C. McPhedran, and Y. S. Kivshar, 2010. Anderson localization of classical waves in weakly scattering metamaterials. Phys. Rev. B 81:075124. http://link.aps.org/doi/10.1103/PhysRevB.81.075124.

12. Kirk, J. T., 1994. Light and photosynthesis in aquatic ecosystems. Cambridge university press.

13. Karpiuk, T., N. Cherroret, K. L. Lee, B. Gremaud, C. A. Müller, and C. Miniatura, 2012. Coherent forward scattering peak induced by anderson localization. Physical review letters 109:190601.

14. Aegerter, C. M., M. Storzer, and G. Maret, 2006. Experimental determination of critical exponents in Anderson localisation of light. EPL (Europhysics Letters) 75:562.

15. Sperling, T., L. Schertel, M. Ackermann, G. J. Aubry, C. M. Aegerter, and G. Maret, 2016. Can 3D light localization be reached in white paint. New Journal of Physics 18:013039. http://stacks.iop.org/1367-2630/18/i=1/a=013039.

16. Noel, M. W., and C. Stroud Jr, 1995. Young’s double-slit interferometry within an atom. Physical review letters 75:1252.

17. Sterligov, V. A., 2005. Scattering and Reflection of Light by Ordered Mesoporous Silica Films. In Frontiers in Optics. Optical Society of America, FThT5.

18. Fiebig, S., C. M. Aegerter, W. B?hrer, M. St?rzer, E. Akkermans, G. Montambaux, and G. Maret, 2008. Conservation of energy in coherent backscattering of light. EPL (Europhysics Letters) 81:64004. http://stacks.iop.org/0295-5075/81/i=6/a=64004.

19. Zimmermann, B., D. Rolles, B. Langer, R. Hentges, M. Braune, S. Cvejanovic, O. Gessner, F. Heiser, S. Korica, T. Lischke, et al., 2008. Localization and loss of coherence in molecular double-slit experiments. Nature Physics 4:649–655.

20. Lagendijk, A., B. van Tiggelen, and D. S. Wiersma, 2009. Fifty years of Anderson localization. Phys. Today 62:24–29.

21. Pradhan, P., and N. Kumar, 1994. Localization of light in coherently amplifying random media. Physical Review B 50:9644.

22. Storzer, M., P. Gross, C. M. Aegerter, and G. Maret, 2006. Observation of the critical regime near Anderson localization of light. Physical Review Letters 96:063904.

23. Watson Jr, G., P. Fleury, and S. McCall, 1987. Searching for photon localization in the time domain. Physical review letters 58:945.

24. Wiersma, D. S., P. Bartolini, A. Lagendijk, and R. Righini, 1997. Localization of light in a disordered medium. Nature 390:671–673.

25. Wolf, P.-E., and G. Maret, 1985. Weak localization and coherent backscattering of photons in disordered media. Physical Review Letters 55:2696.

26. Sperling, T., 2015. The Experimental Search for Anderson Localisation of Light in Three Dimensions. Ph.D. thesis.

27. Mostarda, S., F. Levi, D. Prada-Gracia, F. Mintert, and F. Rao, 2013. Structure-dynamics relationship in coherent transport through disordered systems. Nature communications 4.

28. Lee, K. L., B. Gremaud, C. Miniatura, et al., 2014. Dynamics of localized waves in one-dimensional random potentials: Statistical theory of the coherent forward scattering peak. Physical Review A 90:043605.

29. Ghosh, S., N. Cherroret, B. Gremaud, C. Miniatura, and D. Delande, 2014. Coherent forward scattering in two-dimensional disordered systems. Phys. Rev. A 90:063602. http://link.aps.org/doi/10.1103/PhysRevA.90.063602.

30. Etemad, S., R. Thompson, M. Andrejco, S. John, and F. MacKintosh, 1987. Weak localization of photons: termination of coherent random walks by absorption and confined geometry. Physical review letters 59:1420.

31. Sarovar, M., A. Ishizaki, G. R. Fleming, and K. B. Whaley, 2010. Quantum entanglement in photosynthetic light-harvesting complexes. Nature Physics 6:462–467.

32. Kuhl, M., C. Lassen, and B. B. Jorgensen, 1994. Optical properties of microbial mats: light measurements with fiber-optic microprobes. In Microbial mats, Springer, 149–166.

33. Hamada, F., M. Yokono, E. Hirose, A. Murakami, and S. Akimoto, 2012. Excitation energy relaxation in a symbiotic cyanobacterium, Prochloron didemni, occurring in coral-reef ascidians, and in a free-living cyanobacterium, Prochlorothrix hollandica. Biochimica et Biophysica Acta (BBA) - Bioenergetics 1817:1992–1997. http://www.sciencedirect.com/science/article/pii/S0005272812001934.

34. Kuhl, M., Mhl, R. N. Glud, H. Ploug, and N. B. Ramsing, 1996. MICROENVIRONMENTAL CONTROL OF PHOTOSYNTHESIS AND PHOTOSYNTHESIS-COUPLED RESPIRATION IN AN EPILITHIC CYANOBACTERIAL BIOFILM1. Journal of Phycology 32:799–812.

35. Wiersma, D. S., 2008. The physics and applications of random lasers. Nature physics 4:359–367.

36. Karbasi, S., K. W. Koch, and A. Mafi, 2012. Wavelength Dependence of the Beam Radius in Anderson Localized Optical Fibers. In Frontiers in Optics 2012/Laser Science XXVIII. Optical Society of America, FTh1D.8. http://www.osapublishing.org/abstract.cfm?URI=FiO-2012-FTh1D.8.

37. Bakke, R., R. Kommedal, and S. Kalvenes, 2001. Quantification of biofilm accumulation by an optical approach. Journal of Microbiological Methods 44:13–26.

